# Rare loss-of-function variants in *KMT2F* are associated with schizophrenia and developmental disorders

**DOI:** 10.1101/036384

**Authors:** Tarjinder Singh, Mitja I. Kurki, David Curtis, Shaun M. Purcell, Lucy Crooks, Jeremy McRae, Jaana Suvisaari, Himanshu Chheda, Douglas Blackwood, Gerome Breen, Olli Pietiläinen, Sebastian S. Gerety, Muhammad Ayub, Moira Blyth, Trevor Cole, David Collier, Eve L. Coomber, Nick Craddock, Mark J. Daly, John Danesh, Marta DiForti, Alison Foster, Nelson B. Freimer, Daniel Geschwind, Mandy Johnstone, Shelagh Joss, Georg Kirov, Jarmo Körkkö, Outi Kuismin, Peter Holmans, Christina M. Hultman, Conrad Iyegbe, Jouko Lönnqvist, Minna Männikkö, Steve A. McCarroll, Peter McGuffin, Andrew M. McIntosh, Andrew McQuillin, Jukka S. Moilanen, Carmel Moore, Robin M. Murray, Ruth Newbury-Ecob, Willem Ouwehand, Tiina Paunio, Elena Prigmore, Elliott Rees, David Roberts, Jennifer Sambrook, Pamela Sklar, David St. Clair, Juha Veijola, James T. R. Walters, Hywel Williams, Swedish Schizophrenia Study, INTERVAL Study, DDD Study, UK10K Consortium, Patrick F. Sullivan, Matthew E. Hurles, Michael C. O'Donovan, Aarno Palotie, Michael J. Owen, Jeffrey C. Barrett

## Abstract

Schizophrenia is a common, debilitating psychiatric disorder with a substantial genetic component. By analysing the whole-exome sequences of 4,264 schizophrenia cases, 9,343 controls, and 1,077 parent-proband trios, we identified a genome-wide significant association between rare loss-of-function (LoF) variants in *KMT2F* and risk for schizophrenia. In this dataset, we observed three *de novo* LoF mutations, seven LoF variants in cases, and none in controls (P = 3.3x10^−9^). To search for LoF variants in *KMT2F* in individuals without a known neuropsychiatric diagnosis, we examined the exomes of 45,376 individuals in the ExAC database and found only two heterozygous LoF variants, showing that *KMT2F* is significantly depleted of LoF variants in the general population. Seven of the ten individuals with schizophrenia carrying *KMT2F* LoF variants also had varying degrees of learning difficulties. We further identified four *KMT2F* LoF carriers among 4,281 children with diverse, severe, undiagnosed developmental disorders, and two additional carriers in an independent sample of 5,720 Finnish exomes, both with notable neuropsychiatric phenotypes. Together, our observations show that LoF variants in *KMT2F* cause a range of neurodevelopmental disorders, including schizophrenia. Combined with previous common variant evidence, we more generally implicate epigenetic dysregulation, specifically in the histone H3K4 methylation pathway, as an important mechanism in the pathogenesis of schizophrenia.

## Introduction

Schizophrenia is a common, debilitating psychiatric disorder characterised by positive symptoms (hallucinations, delusions, and disorganisation), and negative symptoms (impaired motivation, reduced spontaneous speech, and social withdrawal). It is associated with cognitive impairment, decreased social and occupational functioning and increased mortality, with a 12 to 15 year reduction in lifespan [1 - 3]. Schizophrenia has a lifetime risk of ∼0.7% and a substantial genetic component, with a sibling recurrence risk ratio of 9.0 and an estimated heritability of up to 81% [4; 5].

The genetic architecture of schizophrenia involves a combination of common, rare, and *de novo* risk variants. At one end of this spectrum, a genome-wide association study of 36,989 cases identified 108 loci containing alleles of individually small effect (median odds ratio = 1.08) [6], while at the other, at least 11 rare, recurrent copy number variants - for example at chromosomes 1q21.1, 15q13.3, and 22q11.2 - individually confer substantial risk for schizophrenia (ORs 2 — 60) [7 - 10]. A recent case-control exome sequencing study demonstrated a burden of rare disruptive variants across a set of 2,546 genes selected based on a variety of biological hypotheses about schizophrenia risk and previous genome-wide screens including GWAS, CNV, and *de novo* mutation studies [11]. This study did not, however, identify any individual schizophrenia risk genes. Parent-proband trio studies have sought to increase power by focusing on *de novo* mutations: the rarity of damaging events makes it possible to observe statistically significant recurrence of mutations in individual genes with smaller sample sizes than would be required in a case-control design. Three such studies in schizophrenia have found suggestive evidence for candidate genes, including *KMT1D* (also known as *EHMT1), DLG2, TAF13* and *KMT2F* (also known as *SETD1A)* [9; 12; 13], but further evidence is needed to firmly establish these as true susceptibility genes.

Two insights have emerged from these early results in schizophrenia. First, genetic risk loci have implicated general biological processes involved in pathogenesis, including histone methylation (common variants) [14], transmission at glutamatergic synapses, and translational regulation by the fragile X mental retardation protein (rare and *de novo* variants) [11; 12]. Second, studies of common and rare variation support a highly polygenic architecture involving hundreds of genes, suggesting that very large sample sizes will be required to convincingly identify individual risk genes. This polygenicity is reminiscent of other neuropsychiatric disorders, such as autism spectrum disorder (ASD), which required many thousands of exome sequences and the integration of *de novo* mutations with case-control burden of rare variants to identify genes at genome-wide significance [15, 16].

## Results

### Case-control analysis of schizophrenia exomes

We sequenced the exomes of 1,887 (1,488 UK and 399 Finnish) individuals with schizophrenia and 7,585 (5,469 UK and 2,116 Finnish) individuals without a known neuropsychiatric diagnosis. We jointly called each case set with its nationality-matched controls, but still observed substantial batch effects from the use of different exome capture reagents used at different time points in the experiment (Supplementary Figure 1). We therefore performed careful quality control (QC) within each set to narrow our analysis to regions with high quality data in all samples, and to remove outlier samples and variants (Online Methods, Supplementary Figure 2), leaving a total of 1,745 cases and 6,789 controls (Figure 1). To increase power for gene discovery, we combined our dataset with exome sequences of 2,519 Swedish schizophrenia cases and 2,554 controls from a previous study [11]. The average number of coding SNPs and indels varied among these three sample sets due to differences in exome capture technology, QC procedures, and sample ancestry, but were closely matched between cases and controls within each set (Supplementary Figures 3, 4, 5). We restricted our analyses to rare variants, stratified by allele frequency (singletons, < 0.1%, and < 0.5%) and function (LoF and damaging missense variants, Online Methods). In total, this joint discovery set consisted of 357,088 damaging missense and 55,955 LoF variants called in 4,264 cases and 9,343 controls (Figure 1).

**Figure 1.**
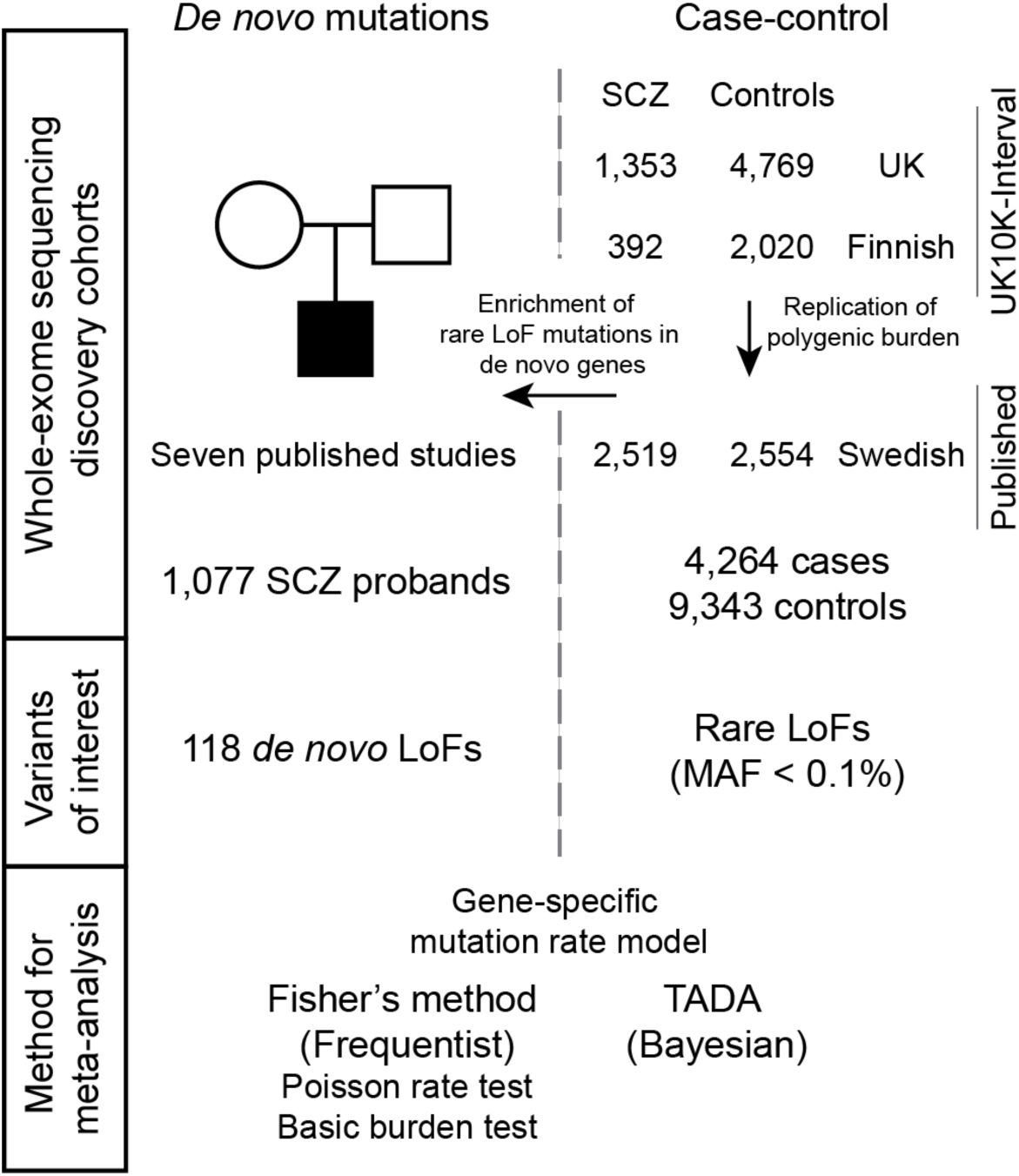
Study design for the schizophrenia exome meta-analysis. The source of sequencing data, sample sizes, variant classes, and analytical methods are described. Details on case-control samples are shown on the right, while parent-proband trios are described on the left.

We replicated the enrichment of rare LoF variants in the previously implicated set of 2,456 genes [11] in our UK and Finnish schizophrenia data sets (μ = 7x10^−4^, Online Methods). Having confirmed that rare disruptive variants spread among many genes are associated with schizophrenia risk, we tested for an excess of disruptive variants within each of 18,271 genes in cases compared to controls (Online Methods). Despite our sample size, the per-gene statistics followed a null distribution in all tests, and we were unable to implicate any gene via case-control burden of disruptive variants (Supplementary Figures 6, 7).

### Loss-of)function variants in *KMT2F*are associated with schizophrenia

To determine whether the integration of *de novo* mutations with case-control burden might succeed in discovering risk genes in schizophrenia, we aggregated, processed, and re-annotated *de novo* mutations in 1,077 schizophrenia probands from seven published studies, and found 118 LoF and 662 missense variants [12; 13; 17 - 21] (Supplementary Table 1). Thirty-eight genes had two or more *de* novo nonsynonymous mutations, two of which *(KMT2F* and *TAF13)* had been previously suggested as candidate schizophrenia genes [12; 13]. We found that the 754 genes with *de novo* mutations were significantly enriched in rare LoF variants in cases compared to controls from our main dataset. The most significant enrichment across allele frequency thresholds and functional class was for the test of LoF variants with MAF < 0.1% (P = 2.1x10^−4^; OR 1.08, 1.02 - 1.14, 95% CI), which we focused on for subsequent analysis.

Motivated by this overlap of genes with *de novo* mutations and excess case-control burden, we meta-analyzed *de novo* variants in the 1,077 published schizophrenia trios with rare LoF variants (MAF < 0.1%) in 4,264 cases and 9,343 controls. We used two analytical approaches, one based on Fisher’s method to combine *de novo* and case-control P-values, and the other using the transmission and *de novo* association (TADA) model to integrate *de novo,* transmitted, and case-control variation using a hierarchical Bayesian framework [15; 22] (Figure 1). We focused on results that were significant in both analyses, and which did not depend on the choice of parameters in TADA (Online Methods). In both methods, loss-of-function mutations in a single gene, *KMT2F* (also known as *SETD1A),* were significantly associated with schizophrenia risk (Table 1a, Fisher’s combined P = 3.3x10^−9^). We observed three *de novo* mutations and seven case LoF variants in our discovery cohort, and none in our controls (Figure 2). In one of the seven case carriers, direct genotyping in parents confirmed that the LoF variant (c.518-2A>G) was a *de novo* event, but genotypes were not available for the other parents. We looked for additional *KMT2F* LoF variants in unpublished whole exomes from 2,435 unrelated schizophrenia cases and 3,685 controls [23], but none were identified (Table 1a). Thus, in more than 20,000 exomes, we observed ten case and zero control LoF variants (corrected OR 35.2, 4.5 - 4528, 95% CI). Although the confidence intervals are wide, rare LoF variants in KMT2F confer substantial risk for schizophrenia. No other gene approached genome-wide significance (Supplementary Table 2, Supplementary Figures 8, 9).

**Table 1a:** Results from statistical tests associating disruptive variants in *KMT2F* to schizophrenia and developmental delay. None of these tests incorporated exomes from the ExAC database. The number of *KMT2F* LoF variants and the sample size of each dataset are indicated in each cell. The statistical tests were performed as follows: 1: a one-sided burden test of case-control LoF variants using Fisher’s exact test, 2: the Poisson probability of observing N *de novo* variants in *KMT2F* given a calibrated baseline gene-specific mutation rate, 3: meta-analysis of *de novo* and case-control burden P-values using Fisher’s combined probability test, 4: the INTERVAL dataset (n = 4,769) were used as matched controls.

| Phenotype | Dataset | *de novo* | Case | Control | Test | P |
| --- | --- | --- | --- | --- | --- | --- |
| Schizophrenia | UK10K-INTERVAL |  | 2/1,353 | 0/4,769 |  |  |
|  | UK10K Finnish |  | 2/392 | 0/2,020 |  |  |
|  | Swedish (published) |  | 3/2,519 | 0/2,554 |  |  |
|  | All case-control |  | 7/4,264 | 0/9,343 | Fisher's exact<sup>1</sup> | 0.0003 |
| | Schizophrenia parent-proband trios | 3/1,077 | | | Poisson exact<sup>2</sup> | $4.6 \times 10^{-7}$ |
| | Case-control + *de novo* (discovery) | 3/1,077 | 7/4,264 | 0/9,343 | Fisher's combined<sup>3</sup> | $3.3 \times 10^{-9}$ |
|  | Swedish (replication) |  | 0/2,435 | 0/3,685 |  |  |
| | All schizophrenia samples | 3/1,077 | 7/6,699 | 0/13,028 | Fisher's combined<sup>3</sup> | $5.6 \times 10^{-9}$ |
| Other neurodevelopmental phenotypes | DDD study | 2/4,281 | 2/4,281 | (see 4) | Fisher's combined<sup>3</sup> | 0.003 |
|  | ASD trios | 0/2,297 |  |  |  |  |
|  | ID trios | 0/151 |  |  |  |  |
| Combined | All samples | 5/7,806 | 9/10,980 | 0/13,028 | Fisher's combined<sup>3</sup> | $3.2 \times 10^{-8}$ |

**Figure 2.**
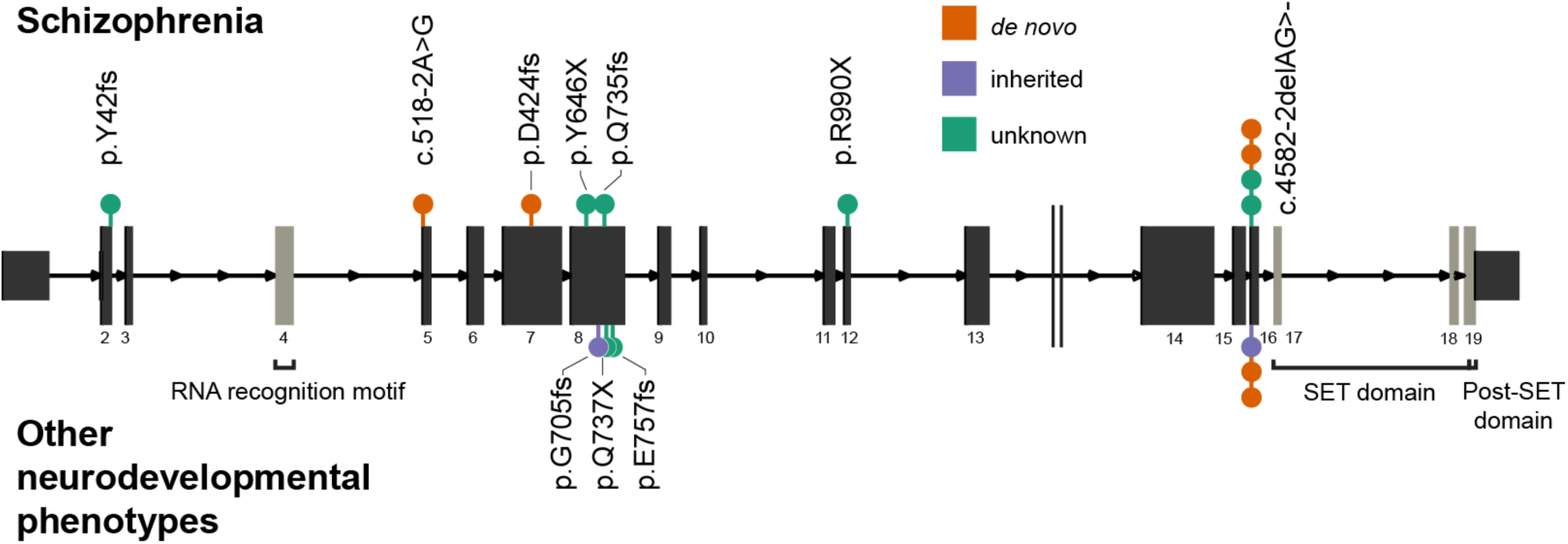
The genomic position and coding consequences of 16 *KMT2F* LoF variants observed in the schizophrenia exome meta-analysis, the DDD study, and the SiSU project. Variants discovered in patients with schizophrenia are plotted above the gene, and those discovered in individuals with other neurodevelopmental disorders (from DDD and SiSU) are plotted below. Each variant is colored according to its mode of inheritance. All LoF variants appear before the conserved SET domain, which is responsible for catalysing methylation. Seven LoF variants occur at the same two-base deletion at the exon 16 splice acceptor (c.4582-2delAG>-).

### Robustness of the *KMT2F*association

To validate our observation of the rarity of disruptive variants in *KMT2F* in unaffected individuals, we examined the exomes of 45,376 individuals without schizophrenia in the Exome Aggregation Consortium (ExAC) database and found only 2 LoF variants [24], which represented a substantial depletion compared to chance expectation (Online Methods, exp. 32.5 LoF SNPs, P = 4.4x10^−8^). *KMT2F* is among the 3% most constrained genes in the human genome [24]; LoF variants in *KMT2F* are almost totally absent in the general population. Four of the ten *KMT2F* carriers with schizophrenia had the same two-base deletion at the exon 16 splice acceptor (c.4582-2delAG>-), at least two of which occurred as *de novo* mutations (Figure 2). Since this variant underpinned the statistical significance of our observation, we investigated it further in several ways. First, to rule out sequencing artifacts, we confirmed a clean call where we had access to the raw sequencing reads (n = 2), and noted that both published *de novo* mutations at this position had been validated with Sanger sequencing [13; 20]. Second, our model, and therefore the test statistic that we report, is dependent on a gene-specific mutation rate (Online Methods). To address the possibility that the recurrent mutation occurs at a hypermutable site (and thus our model is not well calibrated), we determined that our observations would be exome-wide significant (P < 2.5x10^−6^) even if the mutation rate at this position were up to ten-fold higher (7x10^−5^) than the cumulative LoF rate for all other positions in *KMT2F* (6.6x10^−6^). If the two-base deletion mutation rate were truly this high (i.e. greater than 99.99% of all per-gene LoF mutation rates), we would expect to find 6.4 observations in 45,376 non-schizophrenia exomes in ExAC, but we observed only 1 (Fisher’s exact test P = 0.013). Using a minigene construct, we further showed that this two-base deletion resulted in the retention of the upstream intron. This was predicted to lead to the translation of exon 15, the subsequent intron, and an out-of-frame translation of exon 16 resulting in a premature stop codon (Supplementary Figure 11, Online Methods). Finally, if we ignored the *de novo* status of variants in our discovery and replication datasets and used ExAC exomes as additional controls (Online Methods, Table 1b), LoF variants in *KMT2F* were significantly associated with schizophrenia using a basic test of case-control burden (P = 2.6x10^−8^; OR 37.6, 8.0 - 353, 95% CI). Taken together, these analyses excluded many possible artifacts, and provided confidence in our conclusion that LoF variants in *KMT2F* confer substantial risk for schizophrenia.

**Table 1b.**
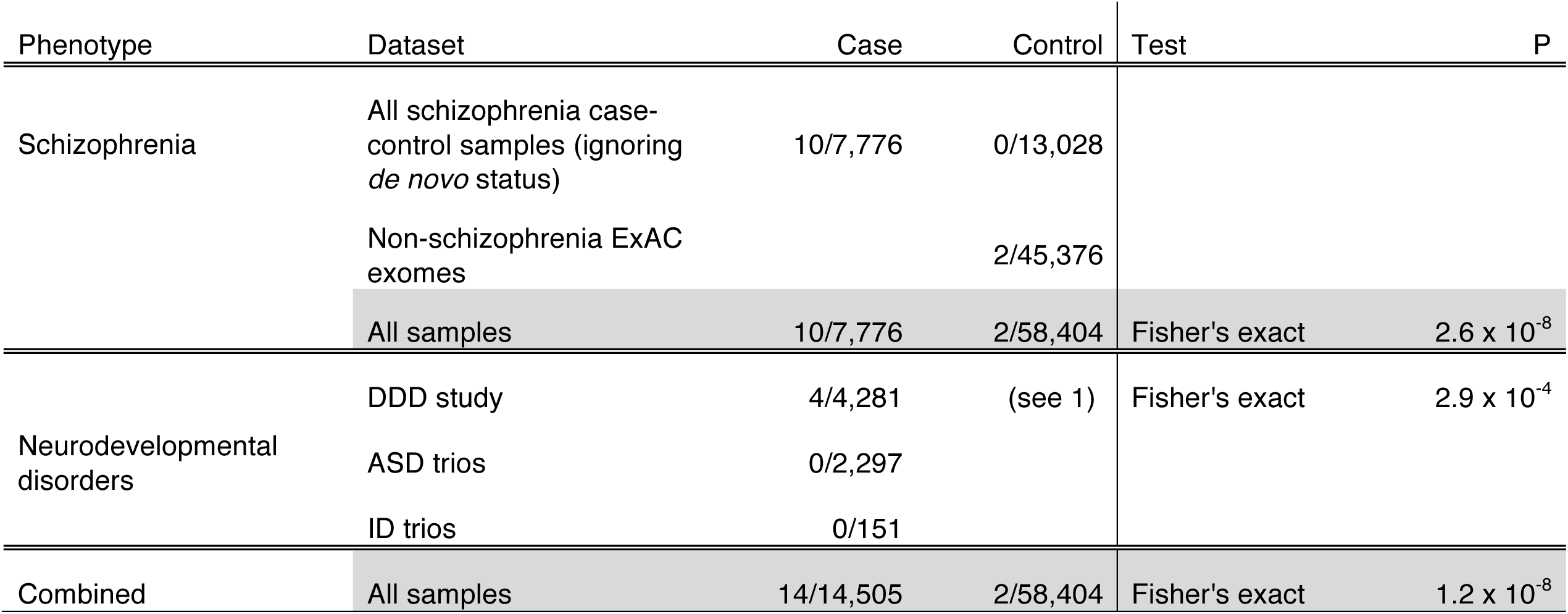
Basic burden tests associating disruptive variants in *KMT2F* to schizophrenia and developmental delay. *De novo* status of variants was ignored and non-schizophrenia exomes from the ExAC database were incorporated as controls. The number of *KMT2F* LoF variants and the sample size of each dataset were indicated in each cell. 1: the full control dataset (n = 58,404) was used to calculate the P-value.

### *KMT2F*LoF variants are associated with developmental disorders

All heterozygous carriers of *KMT2F* LoF variants satisfied the full diagnostic criteria for schizophrenia, including classic positive symptoms such hallucinations, prominent disorganization, and paranoid delusions (Table 2a). Eig patients had evidence of chronic illness, requiring long-term input from psychiatric services. Notably, of the seven *KMT2F* LoF carriers for whom any information on intellectual functioning was available, one was noted to have severe learning difficulties while the six appeared to have mild to moderate learning difficulties. Four patients were noted to have achieved developmental milestones with clinically salient delays (Table 2a). We were unable to confirm if the three Swedish carriers had any form of cognitive impairment. This is consistent with previous reports that individuals with autism or schizophrenia who have *de novo* LoF mutations have a higher rate of cognitive impairment [12; 25].

**Table 2.**
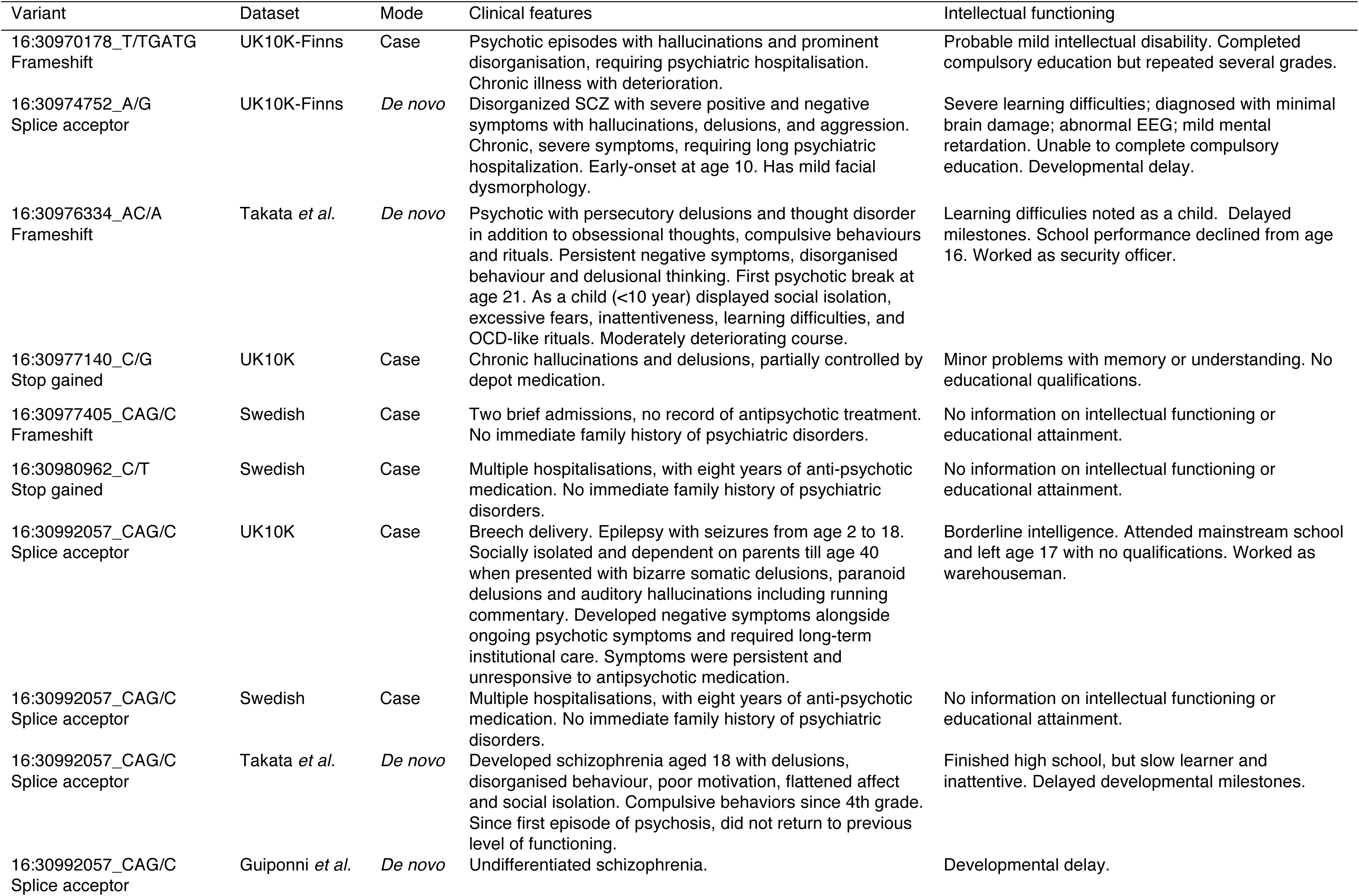
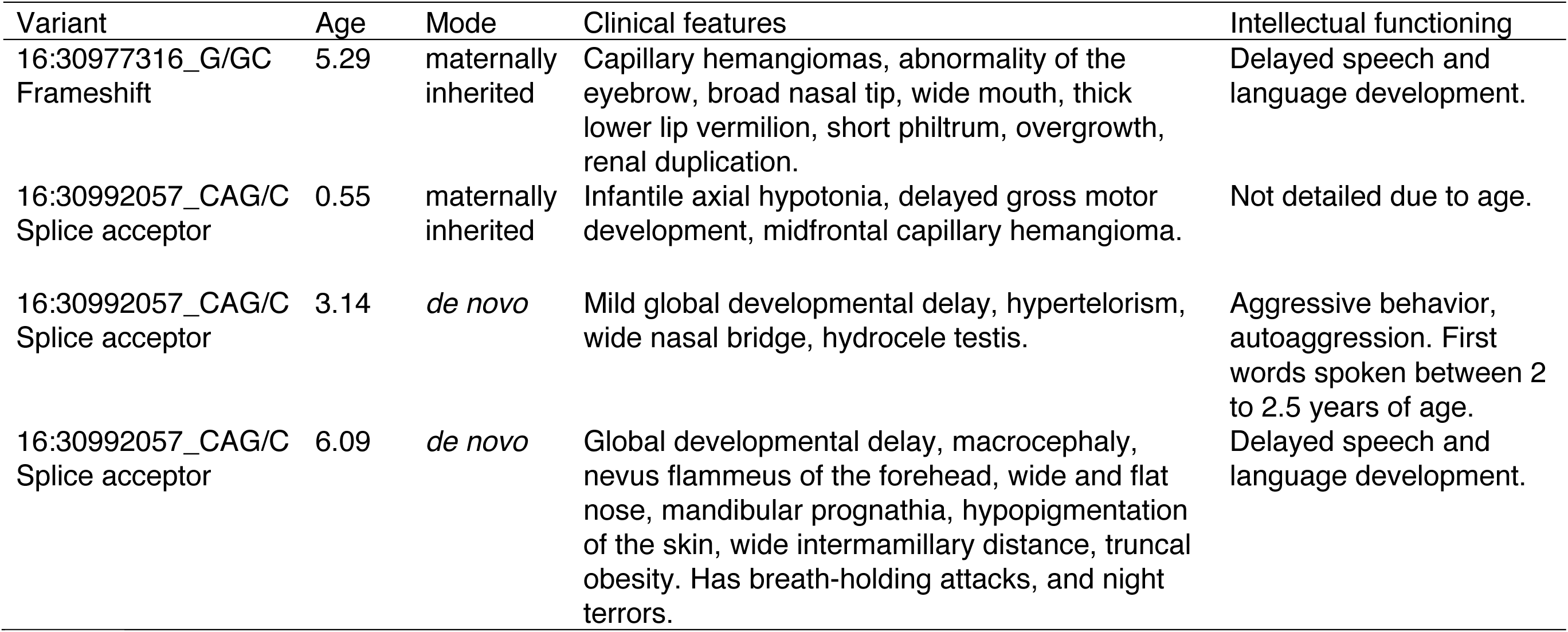
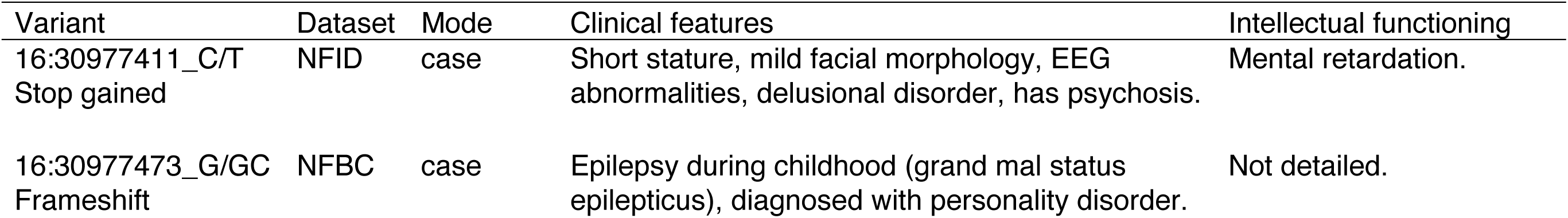
Phenotypes of individuals in the schizophrenia exome meta-analysis (a), the DDD study (b), and the SiSU project (c) who carry LoF variants in *KMT2F.* For each individual, we provide the genomic coordinates of the variant, its mode of inheritance, and the study from which each patient was first recruited. “Clinical features” describes notable neuropsychiatric or neurodevelopmental symptoms in each individual, and “Intellectual functioning” provides additional information on reported cognitive phenotypes. NFID: Northern Finnish Intellectual Disability study; NFBC: Northern Finnish Birth Cohort.

To investigate whether *KMT2F* might play a role in other neurodevelopmental disorders, we looked for *de novo* LoF mutations in *KMT2F* in 3,581 published trios with autism, severe developmental disorders (DD), and/or intellectual disability [15; 26; 27; 28], but found none. We next turned to an additional 3,148 children with diverse, severe, developmental disorders recruited as part of the Deciphering Developmental Disorders (DDD) study, and discovered four probands with LoF variants in *KMT2F* (Table 2b). Three of these were the recurrent exon 16 splice junction indel described above (two *de novo,* one maternally inherited), and the fourth was a maternally inherited frameshift insertion (Figure 2). We validated all four LoF variants using Sanger sequencing. All four probands have developmental delay with additional phenotypes that cluster within the larger DDD study (empirical P = 0.042, Supplementary Information). A fifth proband was found to have a *de novo* 650 kilobase deletion that encompassed *KMT2F* as well as 29 other genes (Supplementary Figure 12, Supplementary Information). *KMT2F* did not reach exome-wide significance as a developmental disorder gene within the DDD study alone (P = 3.0x10^−3^), but when we jointly analyzed all samples, the association was clear to both severe developmental disorders and schizophrenia (P = 3.1X10^−8^, Table 1a). Because all of the DDD *KMT2F* carriers were under 12 years old at recruitment and as schizophrenia rarely manifests at this age [29], it remains unknown if these individuals will develop schizophrenia.

In 5,720 unrelated Finnish individuals exome sequenced as part of the Sequencing Initiative Suomi project (Supplementary Information), we identified two additional heterozygous LoF variants in *KMT2F.* One individual with a stop-gain variant was recruited as part of the Northern Finnish Intellectual Disability cohort with a diagnosis of mental retardation, short stature, mild facial dysmorphology, and EEG abnormalities (Table 2c). Notably, this individual was also diagnosed with delusional disorder and unspecified psychosis at 15 years of age. The second *KMT2F* LoF carrier belonged to the Northern Finnish 1966 Birth Cohort (NFBC), a representative, geographically based population cohort. This individual had epileptic episodes at 7 years of age, and was diagnosed with an unspecified personality disorder by a psychiatrist. Thus, in an additional search for *KMT2F* LoF carriers, only two were found, both in individuals affected by neuropsychiatric disorders.

### Comparison of *de novo* burden between neurodevelopmental disorders

Even though our study had an overall sample size comparable to recent ASD and DD studies that identified 7 ASD genes and 32 DD genes [15; 26], we were only able to implicate a single schizophrenia gene at genome-wide significance. To investigate this further, we aggregated *de novo* mutations identified in 2,297 ASD, 1,113 DD, and 566 control trios with our 1,077 schizophrenia trios, and compared the rates of *de novo* events in each group relative to baseline exome-wide mutation rates (Online Methods). The rates of *de novo* mutations across damaging missense and LoF variants were significantly higher in DD than in ASD, and higher in ASD than in schizophrenia (Figure 3). Indeed, the rate of damaging missense variants in schizophrenia was not different from baseline rates (P = 0.45) and only nominally higher than in controls (P = 0.029), and the rates of LoF variants were only slightly elevated (P = 5.7x10^−3^). In ASD, by contrast, missense (P = 9.4x10^−10^) and LoF (P = 3.7x10^−15^) rates were significantly greater than expectation. In developmental disorders, the rates were even higher (missense: P = 2.5x10^−17^; LoF: P = 1.3x10^−31^) (Figure 3). Across all genes in the genome, the rate of disruptive *de novo* variants differed dramatically across these disorders. Because the recurrence of *de novo* mutations is a particularly powerful way to identify risk genes, the weak excess of *de novo* variants in schizophrenia provides at least a partial explanation for the limited success of this strategy to date in identifying genes for this disorder.

**Figure 3.**
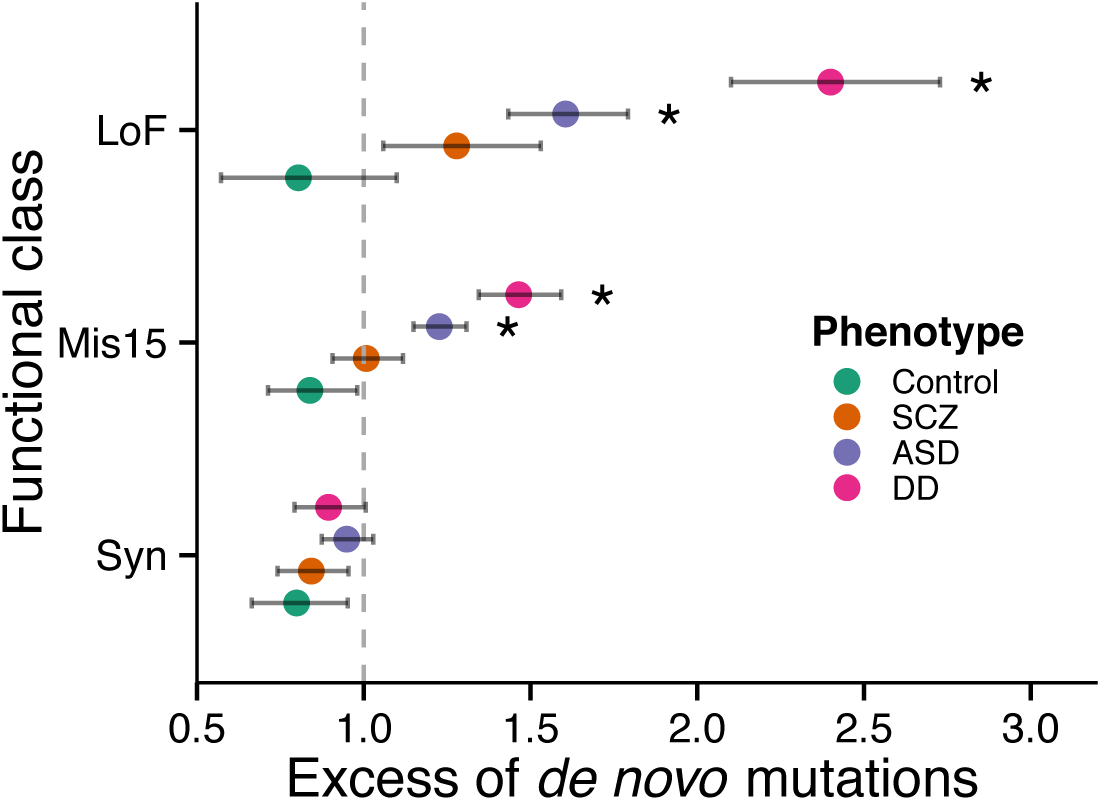
A comparison of genome-wide *de novo* mutation rates in probands with autism, developmental disorders, schizophrenia, and controls. Rates are modeled using calibrated genome-wide mutation rates. Significant excess of *de novo* mutations when compared to the baseline model is marked with an asterisk (P < 4 x 10^−3^, Bonferroni correction for 12 tests). Nominal significance can be inferred from the error bars (95% CI).

## Discussion

In one of the largest exome studies in complex disease to date, we identified a genome-wide significant association to *KMT2F,* the first schizophrenia gene implicated solely by rare coding variants. A previous report [13] suggested *KMT2F* as a candidate schizophrenia gene based on two of the *de novo* mutations included in our analysis, but a much larger sample size was required to conclusively identify it as a risk gene, in keeping with observations in other neurodevelopmental disorder sequencing studies. Indeed, even larger meta-analyses of schizophrenia exomes will be required to rule other candidates in or out, and identify new risk genes. We further demonstrate that LoF variants in *KMT2F* cause a range of severe neurodevelopmental phenotypes in addition to schizophrenia.

*KMT2F* encodes one of the methyltransferases that catalyze the methylation of lysine residues in histone H3. Loss-of-function variants in at least five other genes within this family result in dominant Mendelian disorders characterized by severe developmental phenotypes including intellectual disability [30]. These include Wiedemann-Steiner syndrome (KMT2A), Kleefstra syndrome *(KMT1D/EHMT1),* and Kabuki syndrome *(KMT2D)* (Figure 4). Moreover, rare *de novo* LoF mutations and copy number variants in *KMT2C, KMT2E, KDM5B,* and *KDM6B* have been recently associated with autism risk [16]. The developmental and cognitive phenotypes of *KMT2F* carriers are consistent with these other Mendelian conditions of epigenetic machinery; however, among all genes associated with developmental disorders and intellectual disability, *KMT2F* is the first shown to definitively predispose to schizophrenia, offering insights into the biological differences underlying these conditions [26; 31]. Detailed phenotypes from the DDD study suggest that *KMT2F* carriers may be characterised by more distinctive features, including delayed speech and language development, and facial dysmorphology (Table 2b). While cognitive and developmental phenotypes in our schizophrenia patients are sparser, four individuals had delayed developmental milestones, one is noted as having mild facial dysmorphology and minimal brain damage, and another had epileptic seizures during childhood (Table 2a).

**Figure 4.**
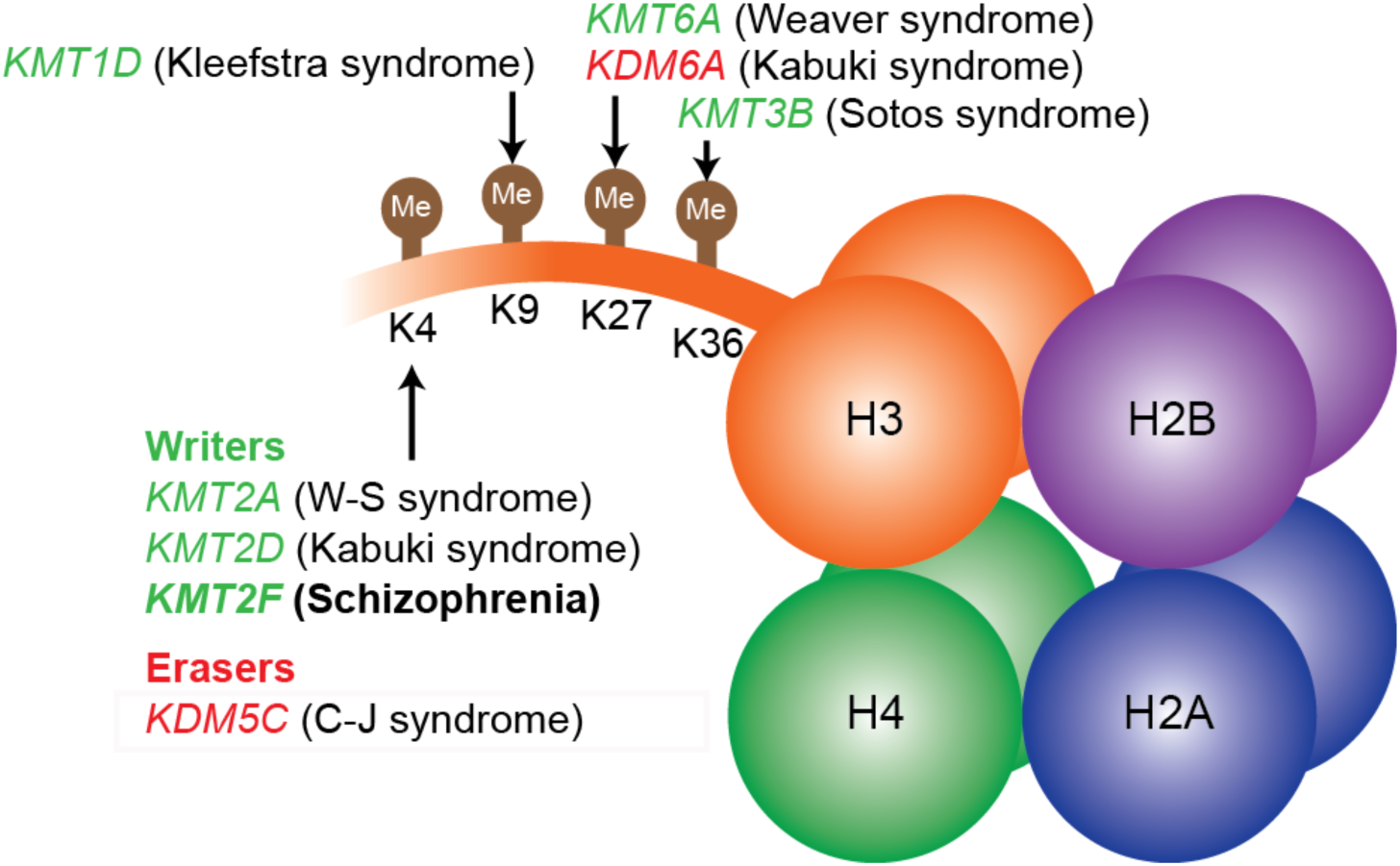
Mendelian disorders of epigenetic machinery at histone H3. Only the tail of histone H3 and its four key lysine residues are illustrated here. Writers (in green) add methyl groups at the specified residue of the histone tail, while erasers (in red) perform targeted demethylation. Disrupting variants in writers and erasers described in the figure result in well-known examples of dominant, highly penetrant disorders characterised by developmental delay and intellectual disability.

The clinical heterogeneity observed in carriers of *KMT2F* LoF variants is reminiscent of at least 11 large copy number variant syndromes (one of which, 16p11.2 is nearby, but not overlapping *KMT2F),* which cause schizophrenia in addition to many other developmental defects [10; 32]. A canonical example is the 22q11.2 deletion syndrome, which is characterised by schizophrenia in 22.6% of adult carriers [33], highly variable intellectual impairment [34], and numerous severe neurological and physical defects [35]. A considerably larger cohort (such as the hundreds of cases of 22q11.2 deletion syndrome studied to date) will be needed to accurately estimate the relative penetrance of *KMT2F* LoF variants for schizophrenia, developmental disorders, and other clinical features.

While disruptions of *KMT2F* are very rare events and occur in only a small fraction of schizophrenia cases (0.13% in our meta-analysis; 0.062% — 0.24% 95% CI), several lines of evidence suggest that histone H3 methylation is more broadly relevant to schizophrenia. The H3K4 methylation gene ontology category (GO:51568) showed the strongest statistical enrichment among 4,939 biological pathways in GWAS data of psychiatric disorders [14]. This category contains 20 genes, including *KMT2F* and six others *(ASH2L, CXXC1, RBBP5, WDR5, DPY30,* and *WDR82)* [36 - 38] that together form the SET1/COMPASS complex, through which *KMT2F* regulates transcription by targeted methylation. Indeed, two of the genes in GO:51568 *(WDR82* and *KMT2E)* are near genome-wide significant associations to schizophrenia [6]. A previous study of *de novo* CNVs in schizophrenia trios identified one deletion and one duplication overlapping *KMT1D* (also known as *EHMT1*), another histone methyltransferase [9] implicated in developmental delay, and a range of congenital abnormalities [39]. Finally, conserved H3K4me3 peaks identified in prefrontal cortical neurons colocalise with genes related to biological mechanisms in schizophrenia including glutamatergic and dopaminergic signaling [40]. Our implication of *KMT2F* therefore contributes to the growing body of evidence that chromatin modification, specifically histone H3 methylation, is an important mechanism in the pathogenesis of schizophrenia.

## Online Methods

### Sample collections

Individuals clinically diagnosed with schizophrenia were recruited and exome sequenced as part of eight neurodevelopmental collections (Aberdeen, Collier, Edinburgh, Gurling, Muir, UK-SCZ, Finnish-SCZ, and Kuusamo) in the UK10K sequencing project. Matched population controls were selected from non-psychiatric arms of the UK10K project, healthy blood donors from the INTERVAL project, and five Finnish population studies (ENGAGE, Familial dyslipidemia, FINRISK, Health 2000, and METSIM). The Swedish schizophrenia case-control and DDD studies had been described previously [11; 26]. The Sequencing Initiative Suomi project consisted a number of prospective and case-control cohorts in Finland. Additional details are in the Supplementary Information.

### Sequence data production

One to three micrograms of DNA was sheared to ∼100 to 400 bp using either a Covaris E210 or LE220 machine (Covaris, Woburn, MA, USA), and processed using Illumina paired-end DNA library preparation. The DNA was enriched using the Agilent SureSelect Human All Exon v.3 or v.5 kits. All libraries were sequenced on the Illumina HiSeq 2000 with 75 base paired-end reads in multiple batches according to the manufacturer’s protocol. Sequencing reads that failed quality control (QC) were first removed using the Illumina GA pipeline. Remaining raw reads were mapped to the reference genome (UK10K: GRCh37, INTERVAL: GRCh37_hs37d5) using BWA (v0.5) [41], and duplicate fragments were marked using Picard (UK10K: v1.36, INTERVAL: v1.114) [42]. We used GATK (UK10K: v1.1-5; INTERVAL: v3.2-2) to perform local realignment around indels, and recalibrate base qualities in each sample BAM [43]. All samples were individually called using GATK Haplotype Caller (v3.2), merged into batches of 200 samples using CombineVCFs, and joint-called using GenotypeVCFs, all at default settings [44; 45]. Due to different captures used in the UK10K and INTERVAL datasets, variant calling was performed at the union of the Agilent v.3 and v.5 captures with 100 base pairs of flanking sequence. To harmonize variant calls across all sequencing batches, we limited subsequent QC and analysis to variants covered at 7x or more in at least 80% of samples in each sequencing batch (Supplementary Figure 1, Supplementary Information).

### Sample-level quality control

Quality control was performed on each population (UK, Finnish, and Swedish) separately. We removed samples with a contamination fraction > 3% estimated using VerifyBamID (v1.0) [46], or low coverage (< 75% of the Gencode v.19 coding region covered at > 10x). Principal components analysis (PCA) was performed using PLINK v1.9 [47] on a set of high-quality (VQSR tranche 99.0%, missingness < 3%, and Hardy-Weinberg P-value < 1x10^−3^), LD-pruned (r^μ^ > 0.2), common (MAF > 5%) SNPs found in our exome capture and in 1000 Genomes Project Phase III data. Ten principal components were estimated using 1000 Genomes samples, onto which we projected all of our cases and controls (Supplementary Figure 5). We verified if samples had the same population ancestry (UK, Finnish or Swedish) as reported in the sample manifests, and excluded individuals who were of non-European ancestry. We estimated kinship coefficients between each sample pair using KING v1.4 [48], and excluded one member of any apparent relative pair (kinship > 0.09375). After sample QC, 6,122 UK samples (1,353 cases and 4,769 controls), 2,412 Finnish samples (392 cases and 2,020 controls), and 5,073 Swedish samples (2,519 cases and 2,554 controls) were available for analysis.

### Variant-level quality control and annotation

We empirically derived thresholds for site and genotype filters that balanced sensitivity and specificity by training on: ExomeChip genotype calls in 295 UK10K cases and doubleton inherited variants (truth sets), and singleton Mendelian inheritance inconsistencies (false set) in 227 trios of the DDD study. We kept SNPs in the VQSR tranche with 99.75% sensitivity and with mean GQ > 30. Individual genotypes were retained if they had a genotype quality (GQ) > 30, alternate allele read depth (DP1) > 2, allelic balance (AB) > 0.2, and AB < 0.8. Using these thresholds, we removed 95.63% of Mendelian errors while retaining 98.38% of doubleton inherited variants, and 99.62% of heterozygous Exomechip SNPs. We kept indels in the VQSR tranche with 99.50% sensitivity, and with mean GQ > 90. Individual genotypes were retained if they had GQ > 90, DP1 > 2, AB > 0.25, and AB < 0.8. Using these thresholds, we removed 92.35% of all indel Mendelian errors, and retained 93.60% of all doubleton inherited indels. We further excluded SNPs and indels with missingness > 20%, Hardy-Weinberg equilibrium *x*^2^ P-values < 1x 10^−8^, variants within low-complexity regions, [49], and indels with more than two alternate alleles, or within 3 base pairs of another indel.

Following sample and variant QC, the per-sample transition-to-transversion ratio was comparable between all populations (mean ∼3.25) (Supplementary Figure 4). We still observed differences in total variant counts among the UK, Finnish, and Swedish collections (Supplementary Figure 3), likely reflecting differences in sequencing depth, capture reagents, sequencing protocol, read alignment, and variant calling. However, variant counts and population genetics metrics were consistent between cases and controls within each population group.

We used the Ensembl Variant Effect Predictor (VEP) version 75 to annotate all variants according to Gencode v.19 coding transcripts [50]. We grouped frameshift, stop gained, splice acceptor and donor variants as loss-of-function (LoF), and missense or initiator codon variants with a CADD Phred score > 15 as damaging missense [51].

### Case-control analysis

To identify genes with a significant burden of rare, damaging variants, we applied the basic burden test, Fisher’s exact test, and the sequence kernel association test (SKAT) as implemented in PLINK/SEQ [52; 53]. For each gene, we tested LoF variants, and LoF combined with damaging missense variants. To evaluate significance, we performed two million case-control permutations within each population (UK, Finnish, and Swedish) to control for ancestry and batch-specific differences. One-sided basic burden and Fisher’s exact tests were applied at three different minor allele frequency (MAF) thresholds (singletons, MAF < 0.1% and MAF < 0.5%). We used default parameters for SKAT (MAF < 5%), and included the first 10 principal components as covariates. Consistent with well-matched cases and controls, we observed no genome-wide inflation in either common or rare variant tests (Supplementary Information, Supplementary Figure 7).

Gene set enrichment analyses broadly followed the methodology described in Purcell *et al.* and implemented in PLINK/SEQ and the SMP utility [11]. The gene set enrichment statistic was calculated as the sum of single gene burden test-statistics corrected for exome-wide differences between cases and controls. Statistical significance was determined through permutation testing as described above. We adopted the min-P procedure to empirically correct for multiple testing: the same order of phenotypic permutations was applied for all tests, and a joint null distribution of minimal P-values was generated to determine the significance of each gene set. The reported odds ratios and confidence intervals from the gene set enrichment analyses were calculated from raw counts without taking into account ancestry and batch-specific differences in cases and controls. Additional details on the methodology and descriptions of the tested gene sets are provided in the Supplementary Information.

### Metaanalysis of *de novo* mutations and case-control burden

Validated *de novo* mutations identified in seven published studies of schizophrenia trios were aggregated for analysis with our case-control cohort (Supplementary Table 1). Recurrence of *de novo* mutations was modeled as the Poisson probability of observing N or more *de novo* variants in a gene given a baseline gene-specific mutation rate obtained from the method described in Samocha *et al.* modified to produce LoF and damaging missense rates for each canonical Gencode v.19 gene [54]. The gene-specific mutation rates in our models have been validated as highly reliable in a previous publication [55], and subsequently used in the main analyses of large-scale exome sequencing of neurodevelopmental disorders with highly replicable results [15,26]. A one-sided Fisher’s exact test was used to model the difference in rare LoF (MAF < 0.1%) burden between cases and controls. Subsequently, *de novo* and case-control burden P-values were meta-analysed using Fisher’s combined probability method with df = 4 (Supplementary Figure 8, 9, Supplementary Table 2). The odds ratios reported were corrected using penalized maximum likelihood logistic regression model (Firth’s method, implemented in the logistf R package).

We also applied the Transmission and Disequilibrium Association (TADA) method as described in He *et al.* [22] and implemented in De Rubeis *et al.* [15]. The robustness of results from TADA depends heavily on the specification of its hyperparameters, which are dependent on the (unknown) genetic architecture of the trait. To ensure our results are robust, we ran the model across a grid of reasonable parameters (Supplementary Information). We found that our signal in *KMT2F* has q-value < 0.01 across a range of plausible parameters (Supplementary Figure 10), but no other gene has q-value < 0.01 under any tested parameterisation (Supplementary Table 3).

### *KMT2F* LoF variants in the ExAC database

We looked in the ExAC database (v0.3) for the LoF variants in *KMT2F.* All exomes were joint-called using the GATK v3.2 pipeline, and included other public exome datasets, such as the 1000 Genomes Project and NHLBI-GO Exome Sequencing Project, with additional quality control compared to their original releases. In 60,706 unrelated exomes, we observed seven LoF variants in *KMT2F.* Since the v0.3 release included the Swedish schizophrenia study, we excluded all samples from this dataset, leaving only four LoF variants in 45,376 exomes without a known neuropsychiatric diagnosis. We next applied the same stringent QC metrics we used in our analysis to ExAC data. We found that the 16:30976302-GC/G indel observed in two individuals was located at the same position as a high-quality SNP, and occurred at a homopolymer run of cytosines. At the genotype level, both calls had a genotype quality (GQ) phred probability of < 40, far lower than used in our study in which we required indels to have a GQ > 90. In addition, the variant has poor allelic balance (AB < 0.15), and the BAM alignment reflected these low-quality metrics (Monkol Lek, personal communication). Given this evidence, we excluded the putative indel. Two high-quality *KMT2F* LoF variants in 45,376 unaffected ExAC exomes remained.

Following the approach in Samocha *et al.,* we determined the significance of the depletion of *KMT2F* LoF variants in ExAC using a signed Z-score of the chi-squared deviation between observed and expected counts [55]. We scaled the expected LoF counts provided by ExAC (43 in 60,706) to 45,376 exomes (exp. 32.5), and calculated the one-tailed P-value of the signed Z-score assuming two observed LoF variants. The degree of constraint relative to other coding genes was based on the pLI score [24].

If we disregarded *de novo* status of our variants, our combined schizophrenia dataset was composed of 7,776 cases and 13,028 controls. After including unaffected ExAC exomes as additional controls, we observed ten LoF variants in 7,776 cases and two LoF variants in 58,404 controls, which was significantly different by a Fisher’s exact test. This result was driven by ten very rare variants in our schizophrenia cases: six observed in only one individual each, and the seventh observed in four individuals. Two of these four were *de novo,* and the other two were found in unrelated individuals of different ancestry (one from Sweden and one from the UK). Similarly, of the two LoF variants in ExAC, one was observed in only one individual and the other was the recurrent indel in an individual of African ancestry. Thus, our burden test of very rare variants in *KMT2F* would not be confounded by systematic differences between subpopulations in the ExAC exomes and our dataset.

### Validation of *KMT2F* variants

We designed primers using Primer3 to produce products between 400 and 600 bp in length centred on the site of interest. Using genomic DNA from all trio members as templates, PCR reactions were carried out using Thermo-Start Taq DNA Polymerase (Thermo Scientific), following the manufacturer’s protocol, and successful PCR products were capillary sequenced. Traces from all trio members were aligned, viewed, and scored for the presence or absence of the variant.

### Functional consequence of the exon 16 splice acceptor deletion

To assess the impact of the exon 16 splice acceptor site variant, we created a custom minigene construct. We cloned the entire 696bp genomic region encompassing exons 15, 16, 17 and intervening introns of human *KMT2F,* fused inframe to a C-terminal GFP. The entire cassette was flanked by a strong upstream promoter and a downstream polyadenylation sequence. Plasmids containing either reference or deletion-containing forms were transfected into HELA cells, which were then grown for 2 days under standard conditions. RNA was extracted (RNEasy, Qiagen) from the transfected cells and used to synthesize cDNA (SuperscriptIII, Invitrogen). We designed minigene-specific primers to avoid amplification of endogenous HELA derived transcripts. The first pair of primers spanned all three exons, thus allowing us to detect overall splicing changes (Pair 1, Forward 2: TCGAAGAGTCATAAACACTGCCATG, Reverse 9: GTGAACAGCTCCTCGCCCTTG). We also designed pairs of exonic, intron-spanning primers to distinguish splicing events upstream (Pair 2, Forward 1: TTTGCAGGATCCCATCGAAGAGTC, exon 16 reverse: CACTGTCCATGATGGCGGAGGTA) and downstream (Pair 3, exon16 forward: CTGCTGAGCGCCATCGGTAC, exon17 reverse: CTGAACTTGTGGCCGTTTACGTC) of exon 16. PCRs were performed on cDNA from two transfection replicates of each sample. Agarose gels identified PCR product size differences (DNA ladder: 2-log ladder, New England Biolabs), which were further analysed by capillary sequencing.

As expected, strong GFP expression was detected from the reference sequence construct. This suggested correct splicing between exons, leading to inframe GFP translation. The mutant form displayed dramatically weaker GFP expression. mRNA was extracted from the transfected cells, and PCRs spanning all three exons revealed an increased transcript size in the mutant form compared to reference (Supplementary Figure 11a). A PCR spanning just the first 2 exons (15/16) revealed a similar shift in size, suggesting that the splice site deletion/mutation was causing intron retention between exons 15 and 16 (Supplementary Figure 11b). Sanger sequencing of the PCR products confirmed this aberrant splicing outcome (Supplementary Figure 11c). The predicted translation product would therefore include translation of exon 15, the subsequent intron, and out of frame translation of exon 16, resulting in a premature stop within this exon. The downstream splicing event to exon 17 was not affected. These data indicate that in human cells, the recurrent indel we observe in probands results in a premature stop codon and a truncated *KMT2F* protein.

### Comparison of *de novo* mutation rates

*De novo* mutations from ASD, DD, and control trios were aggregated from four published studies (Supplementary Information). *De novo* mutations (x_d_) in each neurodevelopmental condition was modelled as x_d_ ∼ Pois(2N_t_[i_G_), where N_t_ is the number of trios, μ_G_ is the genome-wide mutation rate for a particular functional class, and x_d_ is the observed number of *de novo* mutations in N_t_ trios. The genome-wide mutation rate of each variant class was calculated as the sum of all gene-specific mutation rates in Samocha *et al.* [55] (μ_syn_ = 0.137, damaging mis= 0.165, μ_LOF_= 0.043). We modeled *de novo* mutations in control trios to ensure that the genome-wide mutation rates were well calibrated. We reported the probability of observing x_d_ or more mutations in N_t_ trios given the genome-wide mutation rate. We used the Poisson exact test to determine if pairwise differences in *de novo* rates existed between control, SCZ, ASD, and DD trios, and reported the two-sided P-values and rate ratios. Bonferroni correction was used to adjust for multiple testing.

